# Neuroticism Impairs the Use of Reward Values for Decision-Making inMajor Depression

**DOI:** 10.1101/273300

**Authors:** Samuel Rupprechter, Aistis Stankevicius, Quentin Huys, J. Douglas Steele, Peggy Seriès

## Abstract

Depression is a debilitating condition with a high prevalence, but aetiology and pathophysiology are still unclear. Various reward-learning paradigms have been used to show impairments in depression. Both trait pessimism and neuroticism are associated with depression, but their link with the impairments in reward learning and decision-making have not been investigated. A Pavlovian conditioning task was performed by 32 subjects, 15 with depression. Participants had to estimate the probability of some fractal stimuli to be associated with a binary reward, based on a few observations. They then had to make a choice between one of the observed fractals and another target for which the reward probability was explicitly given. Computational modelling was used to succinctly describe participants’ behaviour. Patients performed worse than controls at the task. Computational modelling revealed that this was caused by behavioural impairments during both learning and decision phases. Neuroticism scores across participants were significantly correlated with participants’ inability to follow their internal value estimations. Our results demonstrate behavioural differences in probabilistic reward learning between depressed patients and healthy controls. Neuroticism was associated with the impaired ability to follow internal reward values and consequently with worse decision-making.

## INTRODUCTION

Although major depressive disorder (MDD) is a debilitating condition with a high prevalence and substantial economic impact, aetiology and pathophysiology are still unclear [32]. A core symptom of clinical depression is anhedonia [41] and patients often display impairments in executive function, working memory and attention [27, 34]. It has been proposed that stress blunts reward (reinforcement) learning causing anhedonia [32], and both animal and human research provide evidence supporting this link [8]. Depression and stress are also associated with higher levels of rating scale neuroticism, with the latter being long proposed as a trait risk factor for developing depressive illness [16] and other physical and mental health disorders [40].

Behavioural impairment in MDD has consistently been found with at least two tasks (see Chen and colleagues [13] for a review): the Iowa Gambling Task (see Must and colleagues [29] for a mini review) and the Signal Detection Task (see Huys and colleagues [21] for a meta-analysis). In both paradigms, participants repeatedly choose between options and observe probabilistic reward outcomes based on their choices. Depressed patients are impaired in their ability to properly integrate their reinforcement history to adjust future behaviour. Using the Signal Detection Task, Bogan & Pizzagalli [9] found that, especially in individuals who reported higher levels of anhedonia, experimentally induced stress led to impaired reward processing.

A common symptom during depressive episodes is “bleak and pessimistic views of the future” [41]. The theory of learned helplessness posits that people with a pessimistic explanatory style (attributing their helplessness to a stable, global, internal cause) are at greater risk of developing depression [1]. There exists extensive evidence that patients diagnosed with MDD exhibit features of Beck’s Negative Cognitive Triad, which is characterized by negative and pessimistic views about oneself, the world and the future [6], consistent with a pervasive pessimistic cognitive bias. The Beck Depression Inventory (BDI [5]) and the Beck Hopelessness Scale (BHS [4]) both measure aspects of this triad and Cognitive Behavioural Therapy (CBT), which targets these negative biases can be an effective treatment for depression [3, 12]. Currently, however, the supporting evidence is based almost entirely on subjective clinical interviews and rating scales with little objective behavioural evidence. Here we addressed this issue of subjectivity using a novel experimental paradigm and computational models of decision-making.

We used a probabilistic reward-learning task, which has previously been reported to demonstrate individual behavioural differences that were associated with Life Orientation Test — Revised (LOT-R; measuring optimism) scores [36], as well as neuroticism scores (see Methods and Materials and Supplement) in healthy people. In the task, participants were asked to maximize their rewards by choosing between fractal stimuli, for which they could estimate the probability of reward from previous passive observations, and another target associated with an explicit reward probability value. Here we tested patients with depression as well as healthy controls and used a computational modelling approach to describe their behaviour. This allowed us to formulate specific hypotheses, corresponding to distinct computational models, about both the learning and the decision process during the task and to probe how these processes relate to scores of the rating scales. We were interested in how participants’ ratings of depression severity, optimism and neuroticism affected their performance.

Specifically, we tested whether there was objective evidence for: (a) a behavioural difference in learning and decision-making between MDD subjects and healthy controls, (b) a pessimistic bias about the likelihood of reward in MDD, and (c) a correlation between computational model parameters and ratings of depression severity or neuroticism.

## METHODS AND MATERIALS

### Participants

A total of 39 subjects (Table 1) comprising 19 unmedicated patients meeting DSM-IV criteria for a diagnosis of MDD and 20 control participants without a history of depression or other psychiatric disorder participated. Data collection took place at the Clinical Research Imaging Centre, Ninewells Hospital and Medical School, Dundee. The study was approved by the East of Scotland Research Ethics Service (UK Research Ethics Committee, study reference 13/ES/0043) and all experiments were performed in accordance with relevant guidelines and regulations. Written informed consent was obtained from all subjects.

**Table 1.** Demographics of participants (see Supplement Table S1 for more details). BDI, Beck Depression Inventory; LOTR, Life Orientation Test – Revised; NART, National Adult Reading Test; Data given as n or mean ± std.

| Group | No. Subjects | Age | Sex (F/M) | BDI | Neuroticism | LOT-R | NART |
| --- | --- | --- | --- | --- | --- | --- | --- |
| Patients | 15 | 17 – 41 | 12 / 3 | 24.7 ± 13.1 | 46.3 ± 7.1 | 9.1 ± 5.5 | 46.8 ± 4.2 |
| Controls | 17 | 18 – 33 | 13 / 4 | 4.2 ± 5.6 | 29.8 ± 8.0 | 18.4 ± 3.1 | 46.6 ± 3.2 |

MDD and control groups were matched for age, sex and National Adult Reading Test (NART) scores, which were used to estimate premorbid IQ [11]. Exclusion criteria included claustrophobia, serious physical illness, pre-existing cerebrovascular, neurological disease, previous history of significant head injury, and receipt of any medication likely to affect brain function. Subjects were recruited using the University of Dundee advertisement system HERMES and were paid *£*20 plus up to *£*10 dependent on task performance. Four patients and three controls were excluded from further analysis, after performance results showed that they did not choose the higher reward (in the 48 trials in which the reward probability was not the same) in at least 50% of cases. Two additional participants were excluded from all analysis, because they did not complete the study. This means data was analysed from 15 participants with MDD and 17 controls.

### Experiment

The paradigm (Figure 1) was adapted from Stankevicius and colleagues [36] and performed during fMRI scanning. The experiment was implemented in MATLAB® R2007 (The MathWorks, Inc., Natick, MA) using the Psychophysics Toolbox [10, 31, 25]. Additional details about the experiment are provided in the Supplement and the fMRI analysis will be reported elsewhere. Here we focus on behavioural differences, model fitting and best model identification.

**Figure 1.**
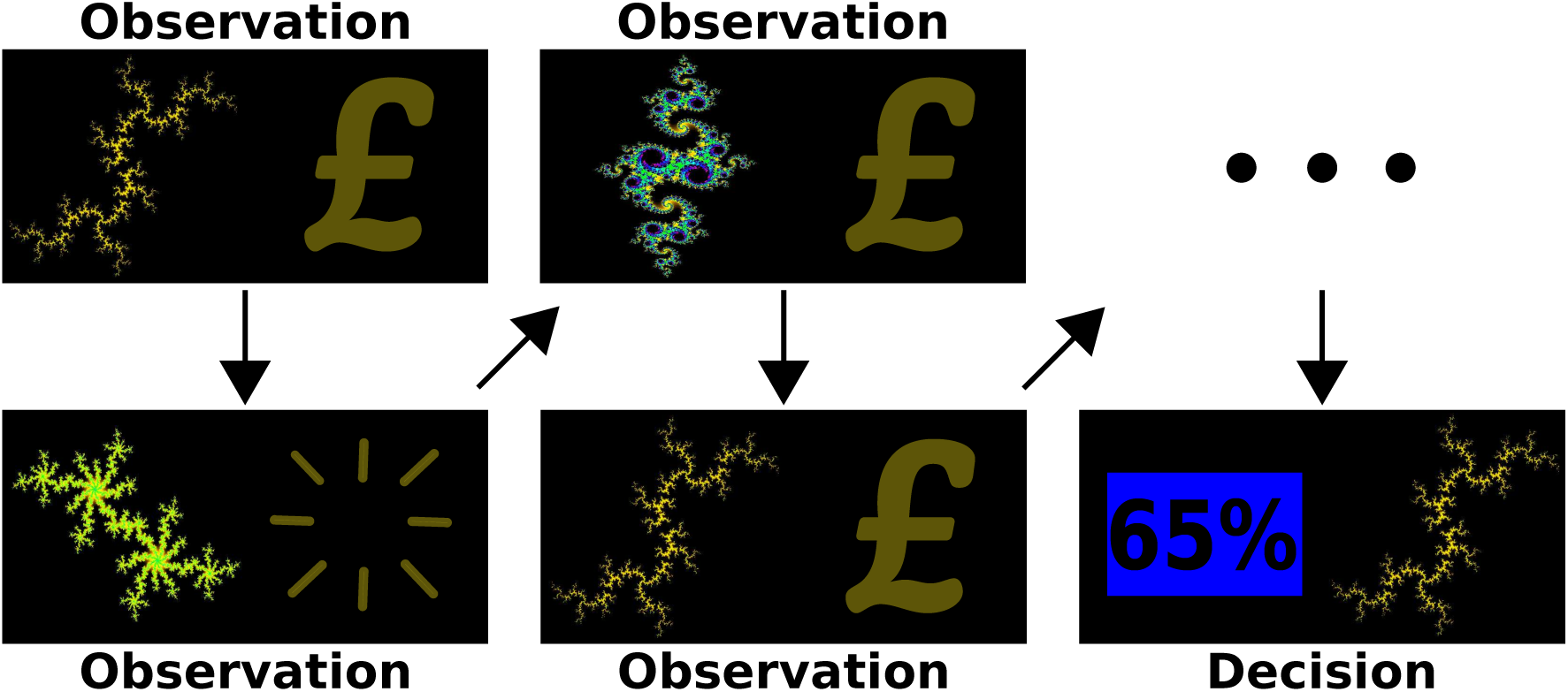
Experimental paradigm. Subject passively observed different fractal stimuli which were followed by reward (a pound symbol) or no reward (blank screen). Interleaved with these observations were decision prompts in which they had to make a choice between one of the observed fractals (for which they could estimate reward probability) and an explicit numeric probability value in order to maximize their reward.

Participants passively observed fractal stimuli, which were followed by either a reward (depicted by a pound symbol) or no reward (no symbol). Interleaved with these observations were decision screens, during which they were asked to make a choice between one of the fractal stimuli they had observed, and a numeric value. Participants were asked to choose the higher probability (or reward) value option, which required them to estimate the value of the fractal stimuli they had observed. There were seven possible differences in the numeric value probability. Either option could have a higher probability value of 10%, 20% or 30% (each of which was the case for 8 decision trials) or they could have the same probability of reward (in 12 trials). Participants observed each fractal four times before a decision screen and each fractal was associated with only a single decision. In total each participant made 60 decisions. The sequence of observations and decisions was pseudo-random, and identical for all subjects. Feedback was only given at the end of the experiment. Data collection for each subject lasted approximately 2 hours, which included collection of rating scale data (see Supplement Table S1).

### Behavioural Performance Data Analysis

We tested for differences in average reaction time, IQ and other questionnaire scores between the groups using Welch’s t tests. Parameters (intercept *α*, slope *β*) of a sigmoid function of the form

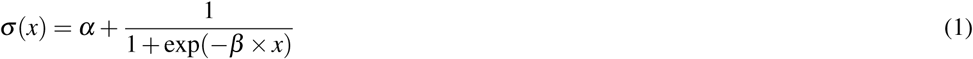

were then fitted to the psychometric curves of individuals to quantify differences in performance as the proportion of fractal stimuli choices as a function of the difference between the reward percentages for the two options.

### Computational Modelling

Three different families of models were fitted to the data (see Table 2 for a summary), representing distinct hypotheses about how participants make decisions during the task. We used formal model selection to choose the most parsimonious model, which was then analysed for differences between MDD and control groups, and for correlations between model parameters and questionnaire scores.

**Table 2.** Model specification.

| Name | V update | p(choose fractal i) | Parameters |
| --- | --- | --- | --- |
| RL-basic | $V_i^{t+1} = V_i^t + \varepsilon(r_i^t - V_i^t)$ | $\sigma(\beta(V_i^t - \phi_i))$ | $v_0, \varepsilon, \beta$ |
| RL-learning | $V_i^{t+1} = V_i^t + \varepsilon^+(1 - V_i^t)r_i^t + \varepsilon^-V_i^t(1 - r_i^t)$ | $\sigma(\beta(V_i^t - \phi_i))$ | $v_0, \varepsilon^+, \varepsilon^-, \beta$ |
| RL-unbiased | $V_i^{t+1} = V_i^t + \varepsilon(r_i^t - V_i^t)$ | $\sigma(\beta(V_i^t - \phi_i))$ | $\varepsilon, \beta$ |
| RL-learning-unbiased | $V_i^{t+1} = V_i^t + \varepsilon^+(1 - V_i^t)r_i^t + \varepsilon^-V_i^t(1 - r_i^t)$ | $\sigma(\beta(V_i^t - \phi_i))$ | $\varepsilon^+, \varepsilon^-, \beta$ |
| Leaky-Bayesian | $V_i^{t+1} = AV_i^t + r_i^t$ | $\sigma(\beta(V_i^t/4 - \phi_i))$ | $A, \beta$ |
| Leaky-Bayesian-ρ | $V_i^{t+1} = AV_i^t + \rho r_i^t$ | $\sigma(\beta(V_i^t/4 - \phi_i))$ | $A, \rho, \beta$ |
| Bayesian | $V_i = \frac{n_i + \alpha}{N_i + \alpha + \beta}$ | $\sigma(\gamma(V_i - \phi_i))$ | $\alpha, \beta, \gamma$ |

First, we fitted variations of a family of reinforcement learning (RL) models that incorporate trial-by-trial prediction errors and a learning rate parameter. Such RL models have been used extensively to describe reward-based learning and much research has gone into understanding the connection between prediction errors and the dopamine system [35]. In two of the models (’RL-basic’ and ‘RL-learning‘), the initial value parameter was allowed to vary between 0 and 1, and could therefore act in a similar way as the mean of the prior belief in the Bayesian model (see below). The other two RL models (’RL-unbiased’ and ‘RL-learning-unbiased‘) kept the bias parameter fixed at 0.5, which corresponded to a prior belief that reward was equally likely from the fractal or the explicit option. Two of these models (’RL-learning’ and ‘RL-learning-unbiased‘) aimed at testing whether learning was different following rewards versus no-rewards by including separate learning parameters for each outcome.

Next, we fitted the winning model of Stankevicius and colleagues [36] (see Table 2), which tests the hypothesis that subjects behave as Bayesian observers during the task. This model assumed that at the decision time for a given fractal, participants estimate the number of times the fractal was followed by a reward (the likelihood) and combine this evidence with a prior belief about the probability of rewards associated with the fractals. Although the observations are not modelled on a trial-by-trial basis, this model assumes that the likelihood is computed by (implicitly) counting, and perfectly remembering, the number of times each fractal is associated with reward. In the original experiment, Stankevicius and colleagues [36] found that the mean of the participants’ prior belief distribution correlated positively with their optimism scores (LOT-R). A more recent analysis of the same data also revealed a negative correlation of the prior mean with neuroticism scores (see Supplement). This means optimists and people scoring low on neuroticism overestimated the reward associated with fractal stimuli and that in this task, optimism and neuroticism acted as a prior belief, biasing performance in situations of uncertainty.

This Bayesian model comes with some limitations. First of all, it does not allow us to distinguish between observation and decision phases, because it ignores individual observation trials. More importantly, the model assumes perfect memory of observations, which is an unrealistic assumption, especially since memory impairments in MDD are exceedingly common [27, 34, 17, 26, 19]. To model memory impairments it is necessary to look at individual trials.

To overcome these limitations, we therefore also fitted two decay RL models (’Leaky-Bayesian’ and ‘Leaky-Bayesian-*ρ*’), which include neither a learning rate nor a prediction error, but which include a discounting factor (also termed a ‘memory’ parameter). This decay-RL rule has consistently been found to produce better model fits than the delta-RL rule when compared using the Bayesian Information Criterion for certain tasks [2]. Internal value estimates are updated after observing fractal *i* and associated reward *r* at observation *t* as

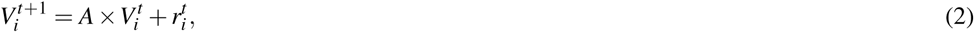

where A (0 < *A* < 1) is the discounting factor (the closer it is to 0, the more a subject “forgets” about their observations and the less they take into account previously observed rewards) and 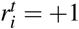 if observation *t* of fractal *i* was rewarded and 0 otherwise. Initial internal values were set to zero. A second model in this family (Leaky-Bayesian-*ρ*) includes a scaling (“reward sensitivity”) parameter on observed rewards, to capture participants’ subjective valuations of observed rewards. Notably, reward processing (dysfunction) has been identified as a promising phenotype of depression [32].

The probability of choosing an action was calculated by passing estimated and explicitly displayed reward probability values through a softmax function. For the Leaky-Bayesian model, fractal *i* was chosen (as opposed to the displayed reward probability *ϕ*_*i*_) with probability

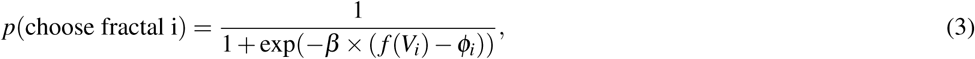

where *f* (*x*) = *x/*4 is a deterministic function which transforms the internal value estimates to a probability comparable to *ϕ*. The shape of the sigmoid function was determined by the *β* parameter. The higher this inverse temperature parameter, the more deterministic decisions become, while lower values lead to “noisier” decision-making. When the values of actions are unknown, this parameter governs the balancing of exploration and exploitation in reinforcement learning [37]. Higher values mean actions are chosen more greedily, lower values lead to suboptimal actions being chosen more often to explore the environment. Here participants were asked to maximize their reward, which means they were asked to always choose the option with the higher probability of reward and there was no advantage of “exploring” the other option. Each fractal was only associated with a single decision and feedback was only given at the end of the experiment and not after each decision. This makes it unlikely that individuals consciously decided to choose the option they thought had a lower probability just to explore the alternative. More plausibly, participants made wrong choices when they either were not certain about what they had observed or had incorrectly estimated the probability of a certain fractal leading to reward. In previous work, the inverse temperature parameter has also been interpreted as reward sensitivity [22] and it has been argued that in some cases it can be substituted exactly for a reward sensitivity parameter [21] in a RL model. This however only holds when all options in the softmax are estimated from observed rewards, which is not the case here. Instead, a high inverse temperature means that participants were able to perform better in the highly uncertain environment and put more trust in their own estimations. Conversely, a lower *β* value meant that participants put less trust in their estimations and chose more randomly.

### Model Fitting and Model Comparison

Model fitting and comparison procedures that have previously been described by Huys and colleagues [20] were used. Parameters were maximum a posteriori (MAP) estimates incorporating an empirical prior, estimated from the data. Parameters were initialized with maximum likelihood values; then an expectation-maximization procedure was used to iteratively update the estimates (see Supplement).

We calculated the integrated Bayesian Information Criterion (iBIC [20]) for all fitted models to find the model that best fitted the data, taking into account complexity. For our winning model we also tested whether our groups were better described using a shared population prior or separate priors for each group.

Simulations were run to verify that both the fitting and comparison procedure recovered reasonable parameters and chose the correct type of model when generating and re-fitting data using known parameters and models (see Supplement).

## RESULTS

### Model-free Analysis

A summary of all questionnaire scores of the two groups is displayed in Supplement Table S1. National Adult Reading Test (NART) scores indicated no difference in IQ between the groups (*p* = .875). Overall, participants did not respond in 17 of 1920 trials (0.89%). Mean response times were not significantly different between groups (RT patients *µ± σ* = 2286 ± 455*ms*; RT controls *µ* ± *σ* = 2185 ± 360*ms*; *t*(26.6) = 0.692, *p* = .495).

Figure 2 shows the fitted sigmoid curves using the average of the fitted parameters for each group. The fitted offset parameter (*α*) was not significantly different between groups (*t*(28.1) = −0.023, *p* = .982), but the slope parameter (*β*) was significantly different (*t*(26.3) = −2.383, *p* = .025), with controls having steeper curves (*β* controls *µ* ± *σ* = 0.566 ± 0.316), indicating they were significantly better at learning (*β* patients *µ* ± *σ* = 0.350 ± 0.185). Notably, the slope parameter was positively correlated with extraversion (*r* = 0.421, *BF* 10 = 2.38, *p* = .016) and negatively correlated with neuroticism (*r* = −0.414, *BF* 10 = 2.16, *p* = .018) scores. Stankevicius and colleagues [36] found a systematic bias in optimistic people towards choosing fractals. We did not find such a systematic bias in healthy participants (as compared to MDD patients) towards choosing fractals, but the difference in the slope parameters indicated performance differences between the groups that we further examined using computational modelling. We were particularly interested in understanding whether those differences stemmed from observation phase or decision phase abnormalities.

**Figure 2.**
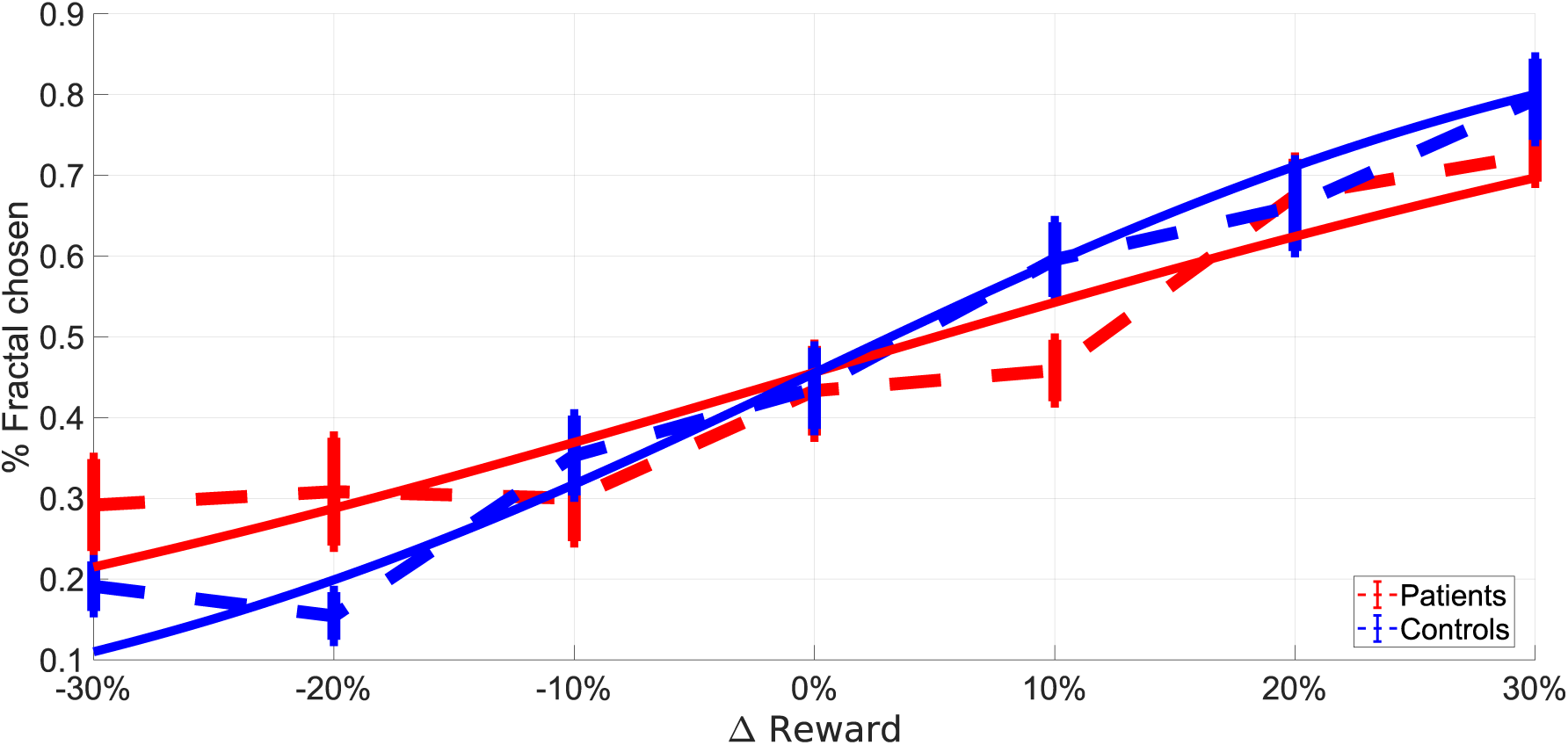
Average sigmoid functions (solid lines) fitted to psychometric curves (dashed lines) of the two groups. Dashed lines depict the average proportion of responses in which the fractal was chosen as a function of the difference between estimated and explicit reward probabilities. Solid lines show the average of simple sigmoid functions fitted to the psychometric curves of individuals. A perfect observer would never choose the fractal when the explicit probability is higher (−30%, −20%, −10%) and always choose the fractal when the estimated probability is higher (10%, 20%, 30%). An unbiased observer would be expected to choose the fractal in half of the trials when reward probability is the same for both options. Error bars represent standard errors.

### Model-based Analysis

Model selection using iBIC showed that the Leaky-Bayesian model best described participants’ performance in our data (Figure 3) and it was best fit using a single population prior (ΔiBIC = 13.5). Figure 4 shows simulations of Leaky-Bayesian using the fitted parameters.

**Figure 3.**
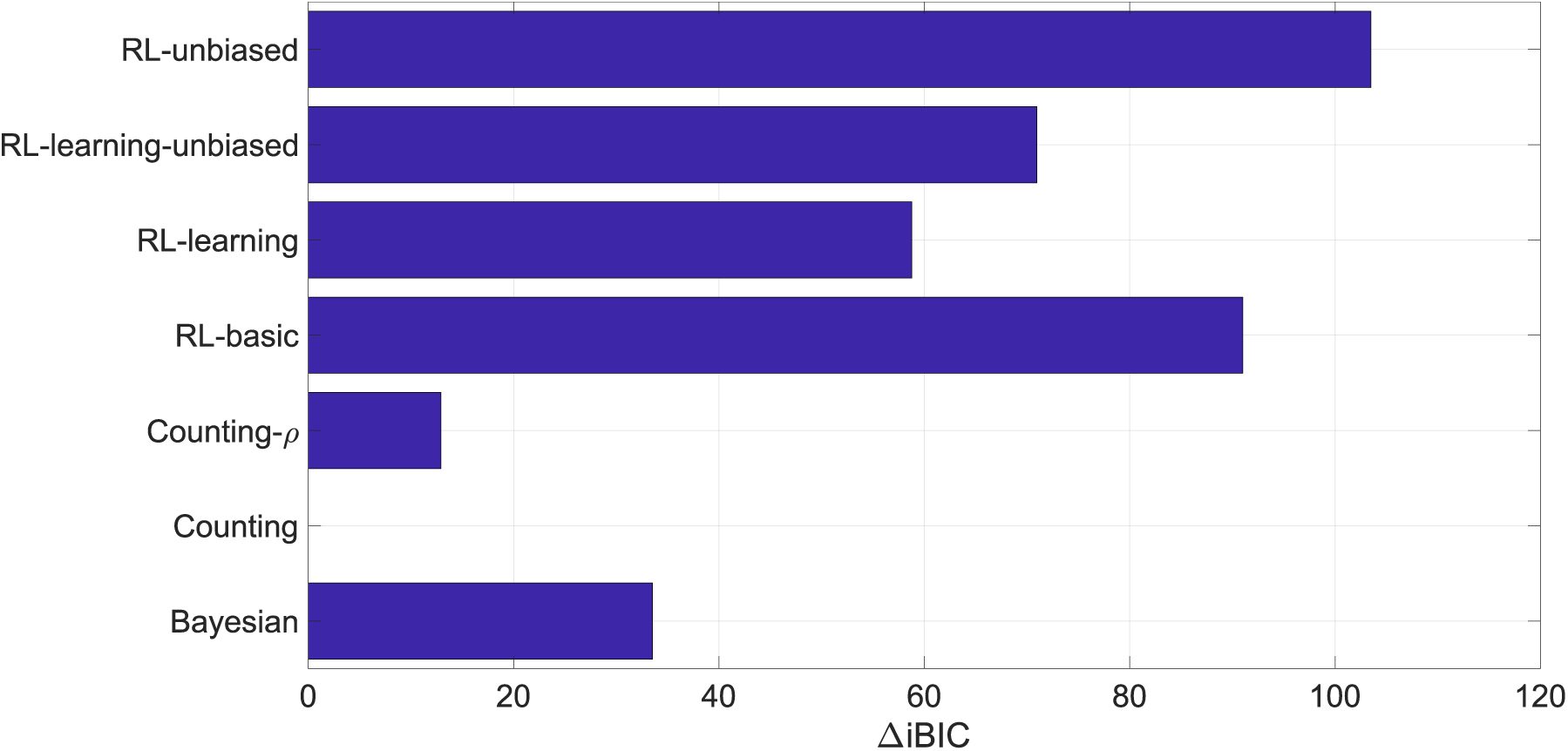
Results of the model comparison. iBIC values of different models relative to the best fitting model Leaky-Bayesian. A difference of 10 or higher is considered strong evidence for the model with the lower value [21].

**Figure 4.**
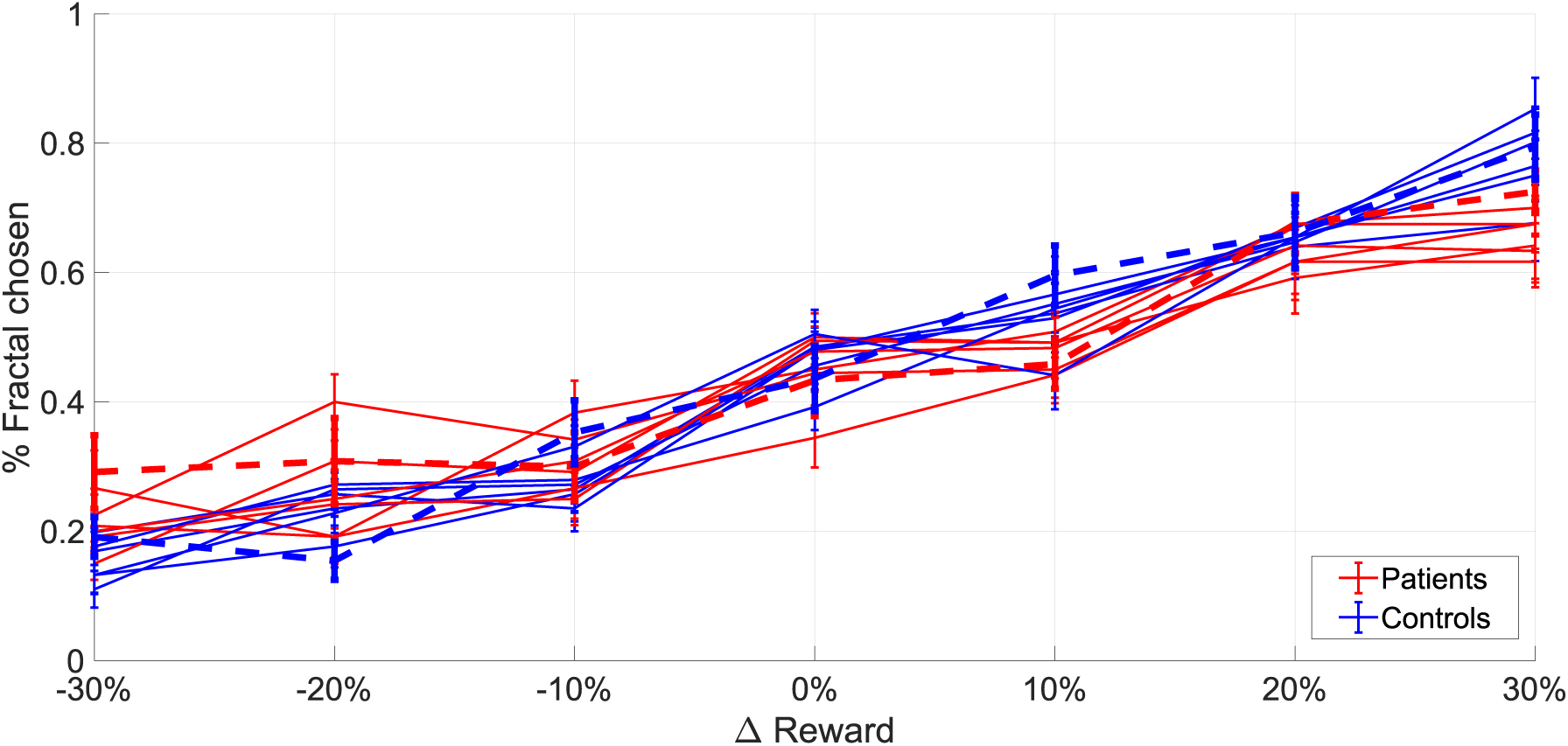
Simulations of the Leaky-Bayesian model. Dashed lines depict the average proportion of responses in which the fractal was chosen as a function of the difference between estimated and explicit reward probabilities. Each solid line also represents this proportion, but for a dataset generated from with best fitting model (Leaky-Bayesian) using the estimated best-fitting parameters. Error bars represent standard errors.

The memory parameter differed significantly between groups (*p* = .031; A patients *µ* ± *σ* = 0.90 ± 0.04, median = 0.91; A controls *µ* ± *σ* = 0.92 ± 0.09, median = 0.96). This indicates that patients discounted their estimated values more than controls on each trial, possibly indicating impairments in working memory. The choice sensitivity parameter (*β*) was also significantly different between groups (*p* = .019; *β* patients *µ* ± *σ* = 4.67 ± 1.45, *β* controls *µ* ± *σ* = 5.89 ± 1.33), meaning that controls found it easier to follow their internal estimations, while patients chose more randomly. There was a trend suggesting a correlation between parameter estimates (*r* = 0.349, *BF* 10 = 0.91, *p* = .051). A negative correlation would mean that variations in one of parameters can account for variations in the other parameter (which could cause problems during parameter estimation), but since this correlation was positive, it is unlikely that this was an issue during inference.

Neuroticism was significantly negatively correlated with the inverse temperature parameter (*r* = −0.491, *BF* 10 = 7.75, *p* = .004), as depicted in Figure 5. This suggests a strong link between participants’ neuroticism and their difficulty in making decisions based on their internal value estimations. High neuroticism was related to much more variable decision-making than low neuroticism.

**Figure 5.**
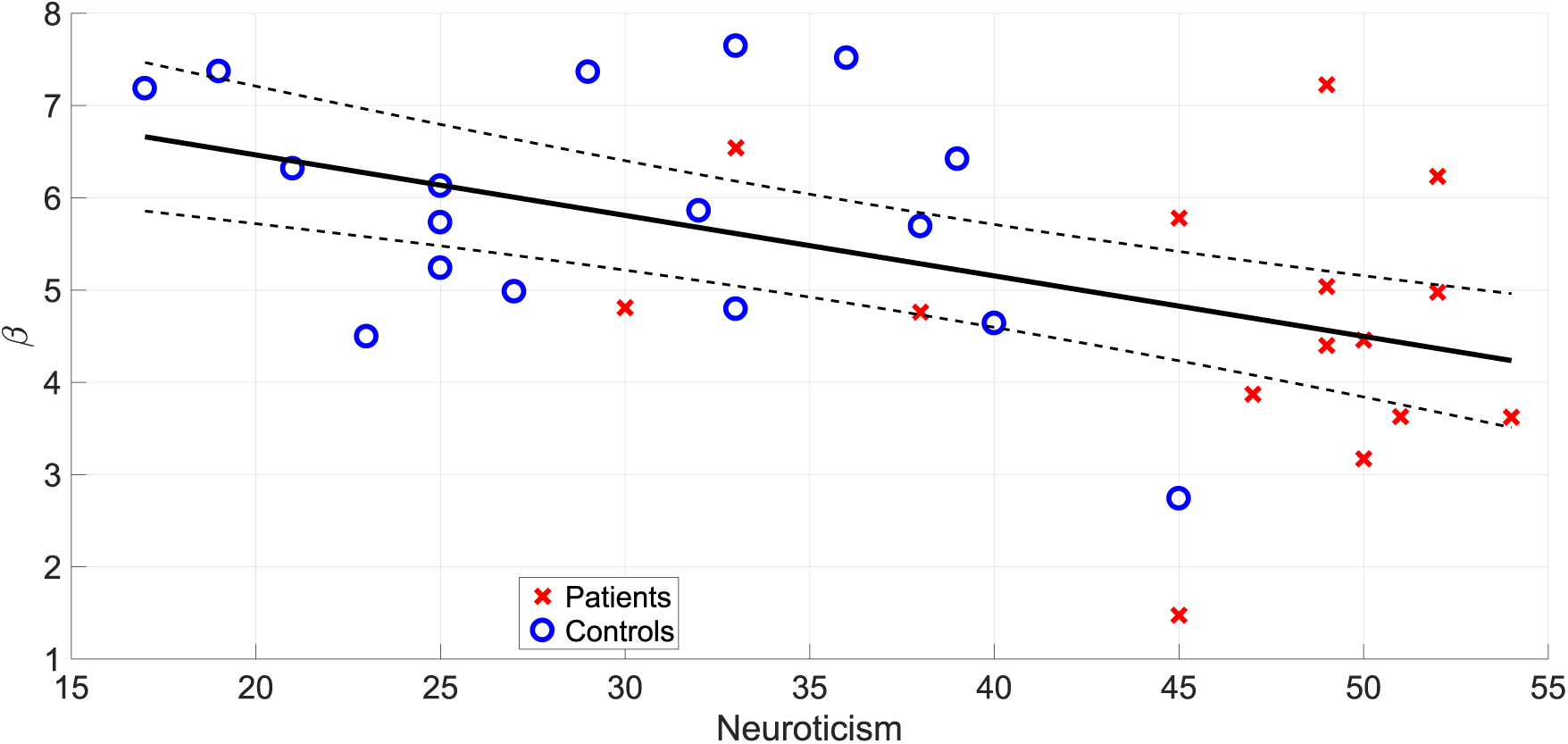
Correlation between neuroticism scores and estimations of the *β* parameter capturing participants’ ability to follow internal value estimations in the best fitting model (Leaky-Bayesian). The lines show the regression line with 95% confidence interval (*r* = −0.491, *BF* 10 = 7.75, *p* = .004).

There were also positive correlations between *β* and extraversion (*r* = 0.423, *BF* 10 = 2.46, *p* = .016), RSE (*r* = 0.410, *BF* 10 = 2.01, *p* = .020), and LOT-R (*r* = 0.382, *BF* 10 = 1.38, *p* = .031) scores, with this being at an “anecdotal” level of evidence (*BF* 10 *<* 3 [39]). Since extraversion, RSE and LOT-R scores were negatively correlated with neuroticism scores (Supplement Figure S1) an additional analysis was done calculating the correlations between those three scores and *β* while controlling for neuroticism scores. None of these partial correlations remained significant (*r* = −0.008, *p* = .968; *r* = −0.124, *p* = .507; *r* = −0.175, *p* = .347; for extraversion, RSE and LOT-R respectively), indicating that the correlation with neuroticism was the driving factor in their correlation with the estimated *β* value.

Further analyses are reported in the Supplement.

## DISCUSSION

Here we used a probabilistic reward-learning task associated with computational modelling to capture behavioural differences between groups of depressed and healthy participants. We found evidence for impairments in MDD subjects during both learning and decision-making. Our results demonstrate a strong inverse relationship between participants’ neuroticism rating and their ability to make decisions based on their internal value estimations. MDD patients also showed decreased memory of observed rewards throughout the task. However, we did not find evidence for a systematic pessimistic bias about the likelihood of reward in depressed participants.

Depression is characterized by behavioural, emotional and cognitive symptoms [19]. It is well established that MDD patients display cognitive impairments including deficits in executive function, working memory, attention and psychomotor processing speed [27, 34]. Behavioural differences in reinforcement learning performance between groups of depressed and healthy participants have been reported previously (see Chen and colleagues [13] for a review). In the Iowa Gambling Task subjects repeatedly choose from one of four different decks of cards with different (unknown to the player) reward and punishment contingencies. High immediate rewards (or losses in an adapted version) are followed by even higher losses (or rewards) at unpredictable points for some decks. Other decks are associated with lower immediate rewards but even lower unpredictable losses. MDD patients typically choose more often from disadvantageous decks, displaying a worsened sensitivity to discriminating reward and punishment (see Must and colleagues [29] for a mini review). In the Signal Detection Task participants observe in each trial one of two hard to distinguish stimuli for a very short time with their task to indicate which stimulus they observed. Correct answers are sometimes rewarded, but unbeknownst to subjects, one of the stimuli is rewarded three times as often as the alternative. Whilst healthy people show a bias towards choosing the more frequently rewarded option, MDD patients do not develop this bias (see Huys and colleagues [21] for a meta-analysis), an effect thought to be related to anhedonia.

In both the Iowa Gambling Task and the Signal Detection Task participants undergo instrumental conditioning, in which chosen actions are reinforced or punished. Subjects learn from their individual choices and the rewards that follow, and will not experience the same reinforcement history, because their rewards depend on their choices. Findings of differences in behaviour or neural activity between groups therefore have to deal with potentially confounding effects of unequal reinforcement histories. Our experiment contains a Pavlovian conditioning phase, during which conditioned stimuli (fractals) are paired with reward and no choices are made. All participants passively observed the exact same sequence of stimuli and these rewards. It is very unlikely that participants learned from their instrumental choices in our task, because each fractal stimulus was only associated with a single decision and feedback was only displayed at the end of the experiment.

Computational modelling was used to capture the behaviour of participants during the task and formal model comparison to choose the best fitting model, from which we identified the best fitting parameters for each participant. MDD patients performed worse on our task and the model-based analysis showed that this was due to differences in two model parameters. First, patients discounted (or forgot) previous reward history more than comparison subjects, consistent with reported impairments in working memory and attentional deficits [27, 34]. Dombrovski and colleagues found suicide attempters (but intriguingly not non-suicidal depressed elderly people) had lower memory parameter values than control participants in a probabilistic reversal learning task [15]. Our finding is also consistent with another recent study by Pulcu and colleagues which reported increased discounting of rewards in MDD [33], although discounting in our task was related to past rewards, while Pulcu and colleagues’ task involved future rewards. Notably, a link between working memory and delay discounting has previously been reported [7, 38]. Second, we also found MDD patients had more difficulty following their internal value estimations of different stimuli, making decisions more randomly. It is possible that patients had a lower confidence in their ability to perform the task, similar to how learned helplessness theories view depression as a consequence of an organism’s diminished belief about its ability to influence outcomes [1]. Taken together, our results therefore suggest that MDD is associated with dysfunctions in both learning and decision-making.

Depression is characterized by dysfunctional processing of rewards during reinforcement learning [18] and there is considerable evidence for a prominent role of stress and stress sensitivity in depression onset; depression could emerge from stress-induced anhedonia [32, 8]. Stress reactivity is thought to be a core aspect of neuroticism, with individuals scoring highly on neuroticism showing greater sensitivity to aversive (stressful) events [30]. Neuroticism is associated with a vulnerability to many common psychiatric disorders including depression [40, 30]. On the basis of animal models of human mood disorder, Deakin and Graeff [14] proposed that serotonin plays a crucial role in resilience, describing how anxiety disorders and depression could arise from impairments in serotonergic pathways. A recent clinical fMRI study [23] strongly supports for Deakin and Graeff’s predictions, and a large population based study concluded that neuroticism increases vulnerability to depression because of increased sensitivity to stressful life events [24].

We found a strong negative correlation between self-reported neuroticism and a model parameter capturing a subject’s ability to use internal value estimates, meaning higher neuroticism scores were associated with a more variable decision process. Taking into account the above literature, a possible explanation for this is that participants with higher stress sensitivity, as indicated by a neuroticism score, were more likely to be depressed, tended to value recent observations of rewards more, had less confidence in their own estimations and consequently performed worse on the task. Considering the environment in which the task was performed, it is possible that the fMRI scanner acted as a stressor during the study [28], especially impacting more stress-sensitive subjects.

Our sample size of 32 analysed participants was relatively small, which might have influenced the model comparison procedure and comparison with our previous study [36]. Nevertheless, model recovery simulations (see Supplement) indicated that all models, with the exception of Leaky-Bayesian-*ρ*, could reliably be distinguished from our Leaky-Bayesian model. There is a chance that Leaky-Bayesian won over Leaky-Bayesian-*ρ* mainly because we did not have enough data and it will be useful for future research to include more participants to assess whether an additional reward sensitivity parameter is useful in describing differences in behaviour during the task. As always in computational modelling studies, there is also a possibility that there exists an untested model that better describes the choices of participants. The task was quite difficult and participants had to be able to deal with large uncertainties about their estimations. Future research could add some easier options with a greater difference in probabilities to get a better baseline of participants’ decision-making.

In conclusion, our results demonstrate impairments in MDD in a probabilistic reward-learning task during both learning and decision-making phases of the experiment. A strong negative correlation was found between neuroticism ratings and subjects’ abilities to follow internal value estimations. High neuroticism was primarily found in MDD patients. Results are consistent with the interpretation that participants with high neuroticism, an indicator for high stress sensitivity, had a diminished ability (possibly due to decreased confidence) to correctly estimate reward probabilities and consequently displayed worse decision-making.

## ACKNOWLEDGEMENTS

SR received a Principal’s Career Development Ph.D. Scholarship from the University of Edinburgh. AS and data collection were supported by grants EP/F500385/1 and BB/F529254/1 for the University of Edinburgh School of Informatics Doctoral Training Centre in Neuroinformatics and Computational Neuroscience (http://www.anc.ed.ac.uk/dtc/) from the UK Engineering and Physical Sciences Research Council (EPSRC), UK Biotechnology and Biological Sciences Research Council (BBSRC), and the UK Medical Research Council (MRC).

We thank Frank Karvelis, James Raymond and Aleks Stolycin for valuable feedback on previous versions of this manuscript.

## AUTHOR CONTRIBUTIONS STATEMENT

A.S., Q.H., D.S., and P.S. conceived the experiment. A.S. and D.S. collected the data. S.R. analysed the data. All authors wrote the manuscript.

## ADDITIONAL INFORMATION

### Competing financial interests

S.R., A.S., Q.H. and P.S. reported no biomedical financial interests or potential conflicts of interest.

D.S. is currently receiving financial support from Indivior for a different study on subjects with opiate dependency.

## REFERENCES

[1] Lyn Y Abramson, Martin E Seligman, and John D Teasdale. Learned helplessness in humans: Critique and reformulation. Journal of abnormal psychology, 87(1):49, 1978.

[2] Woo-Young Ahn, Adam Krawitz, Woojae Kim, Jerome R Busemeyer, and Joshua W Brown. A model-based fmri analysis with hierarchical Bayesian parameter estimation. Journal of neuroscience, psychology, and economics, 4(2):95, 2011.

[3] Aaron T Beck. The current state of cognitive therapy: a 40-year retrospective. Archives of General Psychiatry, 62(9):953–959, 2005.

[4] Aaron T Beck and Robert A Steer. Beck Hopelessness Scale. Psychological Corporation San Antonio, TX, 1988.

[5] Aaron T Beck, Calvin H Ward, Mock Mendelson, Jeremiah Mock, and John Erbaugh. An inventory for measuring depression. Archives of general psychiatry, 4(6):561–571, 1961.

[6] A.T. Beck, A.J. Rush, B.F. Shaw, and G. Emery. Cognitive Therapy of Depression. Guilford clinical psychology and psychotherapy series. Guilford Press, 1979.

[7] Warren K Bickel, Richard Yi, Reid D Landes, Paul F Hill, and Carole Baxter. Remember the future: working memory training decreases delay discounting among stimulant addicts. Biological psychiatry, 69(3):260–265, 2011.

[8] Ryan Bogdan, Yuliya S Nikolova, and Diego A Pizzagalli. Neurogenetics of depression: a focus on reward processing and stress sensitivity. Neurobiology of disease, 52:12–23, 2013.

[9] Ryan Bogdan and Diego A Pizzagalli. Acute stress reduces reward responsiveness: implications for depression. Biological psychiatry, 60(10):1147–1154, 2006.

[10] David H Brainard. The psychophysics toolbox. Spatial vision, 10:433–436, 1997.

[11] Peter Bright, ELI Jaldow, and Michael D Kopelman. The national adult reading test as a measure of premorbid intelligence: a comparison with estimates derived from demographic variables. Journal of the International Neuropsychological Society, 8(06):847–854, 2002.

[12] Andrew C Butler, Jason E Chapman, Evan M Forman, and Aaron T Beck. The empirical status of cognitive-behavioral therapy: a review of meta-analyses. Clinical psychology review, 26(1):17–31, 2006.

[13] Chong Chen, Taiki Takahashi, Shin Nakagawa, Takeshi Inoue, and Ichiro Kusumi. Reinforcement learning in depression: a review of computational research. Neuroscience & Biobehavioral Reviews, 55:247–267, 2015.

[14] JF William Deakin and Frederico G Graeff. 5-ht and mechanisms of defence. Journal of psychopharmacology, 5(4):305–315, 1991.

[15] Alexandre Y Dombrovski, Luke Clark, Greg J Siegle, Meryl A Butters, Naho Ichikawa, Barbara J Sahakian, and Katalin Szanto. Reward/punishment reversal learning in older suicide attempters. American Journal of Psychiatry, 167(6):699–707, 2010.

[16] Conor Duggan, Pak Sham, Alan Lee, Carrine Minne, and Robin Murray. Neuroticism: a vulnerability marker for depression evidence from a family study. Journal of affective disorders, 35(3):139–143, 1995.

[17] Klaus P Ebmeier, Claire Donaghey, and J Douglas Steele. Recent developments and current controversies in depression. The Lancet, 367(9505):153–167, 2006.

[18] Neir Eshel and Jonathan P Roiser. Reward and punishment processing in depression. Biological psychiatry, 68(2):118–124, 2010.

[19] Ian H Gotlib and Jutta Joormann. Cognition and depression: current status and future directions. Annual review of clinical psychology, 6:285–312, 2010.

[20] Quentin JM Huys, Roshan Cools, Martin GÖlzer, Eva Friedel, Andreas Heinz, Raymond J Dolan, and Peter Dayan. Disentangling the roles of approach, activation and valence in instrumental and Pavlovian responding. PLoS Comput Biol, 7(4):e1002028, 2011.

[21] Quentin JM Huys, Diego A Pizzagalli, Ryan Bogdan, and Peter Dayan. Mapping anhedonia onto reinforcement learning: a behavioural meta-analysis. Biology of mood & anxiety disorders, 3(1):12, 2013.

[22] Quentin JM Huys, Joshua T Vogelstein, Peter Dayan, and L Bottou. Psychiatry: Insights into depression through normative decision-making models. In NIPS, pages 729–736, 2008.

[23] Blair A Johnston, Serenella Tolomeo, Victoria Gradin, David Christmas, Keith Matthews, and J Douglas Steele. Failure of hippocampal deactivation during loss events in treatment-resistant depression. Brain, 138(9):2766–2776, 2015.

[24] Kenneth S Kendler, Jonathan Kuhn, and Carol A Prescott. The interrelationship of neuroticism, sex, and stressful life events in the prediction of episodes of major depression. American Journal of Psychiatry, 161(4):631–636, 2004.

[25] Mario Kleiner, David Brainard, Denis Pelli, Allen Ingling, Richard Murray, Christopher Broussard, et al. What’s new in psychtoolbox-3. Perception, 36(14):1, 2007.

[26] Lisa M McDermott and Klaus P Ebmeier. A meta-analysis of depression severity and cognitive function. Journal of affective disorders, 119(1):1–8, 2009.

[27] Roger S McIntyre, Danielle S Cha, Joanna K Soczynska, Hanna O Woldeyohannes, Laura Ashley Gallaugher, Paul Kudlow, Mohammad Alsuwaidan, and Anusha Baskaran. Cognitive deficits and functional outcomes in major depressive disorder: determinants, substrates, and treatment interventions. Depression and anxiety, 30(6):515–527, 2013.

[28] Markus Muehlhan, Ulrike Lueken, Hans-Ulrich Wittchen, and Clemens Kirschbaum. The scanner as a stressor: evidence from subjective and neuroendocrine stress parameters in the time course of a functional magnetic resonance imaging session. International Journal of Psychophysiology, 79(2):118–126, 2011.

[29] Anita Must, Szatmar Horvath, Viola Luca Nemeth, and Zoltan Janka. The Iowa gambling task in depression–what have we learned about sub-optimal decision-making strategies? Frontiers in psychology, 4:732, 2013.

[30] Johan Ormel, A Bastiaansen, Harriëtte Riese, Elisabeth H Bos, Michelle Servaas, Mark Ellenbogen, Judith GM Rosmalen, and André Aleman. The biological and psychological basis of neuroticism: current status and future directions. Neuroscience & Biobehavioral Reviews, 37(1):59–72, 2013.

[31] Denis G Pelli. The videotoolbox software for visual psychophysics: Transforming numbers into movies. Spatial vision, 10(4):437–442, 1997.

[32] Diego A Pizzagalli. Depression, stress, and anhedonia: toward a synthesis and integrated model. Annual review of clinical psychology, 10:393, 2014.

[33] E Pulcu, PD Trotter, EJ Thomas, M McFarquhar, Gabriella Juhász, BJ Sahakian, JFW Deakin, R Zahn, IM Anderson, and R Elliott. Temporal discounting in major depressive disorder. Psychological medicine, 44(09):1825–1834, 2014.

[34] PL Rock, JP Roiser, WJ Riedel, and AD Blackwell. Cognitive impairment in depression: a systematic review and meta-analysis. Psychological Medicine, 44(10):2029, 2014.

[35] Wolfram Schultz. Getting formal with dopamine and reward. Neuron, 36(2):241–263, 2002.

[36] Aistis Stankevicius, Quentin JM Huys, Aditi Kalra, and Peggy Seriès. Optimism as a prior belief about the probability of future reward. PLoS Comput Biol, 10(5):e1003605, 2014.

[37] Richard S Sutton and Andrew G Barto. Reinforcement learning: An introduction, volume 1. MIT press Cambridge, 1998.

[38] Michael J Wesley and Warren K Bickel. Remember the future ii: meta-analyses and functional overlap of working memory and delay discounting. Biological psychiatry, 75(6):435–448, 2014.

[39] Ruud Wetzels and Eric-Jan Wagenmakers. A default Bayesian hypothesis test for correlations and partial correlations. Psychonomic bulletin & review, 19(6):1057–1064, 2012.

[40] Thomas A Widiger and Joshua R Oltmanns. Neuroticism is a fundamental domain of personality with enormous public health implications. World Psychiatry, 16(2):144–145, 2017.

[41] World Health Organization. The ICD-10 classification of mental and behavioural disorders: clinical descriptions and diagnostic guidelines. Geneva: World Health Organization, 1992.

